# Alström Syndrome protein Alms1 is required for maintaining meiotic centriole engagement in spermatocytes of *Drosophila*

**DOI:** 10.1101/2023.09.12.557481

**Authors:** Yanan Hou, Zhimao Wu, Qing Wei

## Abstract

Maintaining proper mother-daughter centriole engagement is crucial for controlling centrosome numbers and ensuring accurate cell division in both mitosis and meiosis. However, the mechanism responsible for this maintenance remains unclear. Here, we report that the *Drosophila* homolog of human ciliopathy gene Alström Syndrome 1 (Alms1) is required for maintaining centriole engagement during spermatocyte meiosis. We demonstrated that deletion of Alms1 results in premature centriole disengagement in prophase of meiosis I, leading to the formation of multipolar spindles and abnormal cell division in *Drosophila* spermatocytes. Further studies reveal that Alms1 localizes to the proximal centrioles, and functions downstream of cartwheel protein Sas-6 to regulate centriole engagement, and its highly conserved ALMS motif is required for Alms1’s function in centriole engagement. Moreover, we show that the centriole size and pericentriolar materials (PCM) have antagonistic roles on maintaining centriole engagement in *alms1* mutant flies. Our findings highlight the critical role of Alms1 as a “glue protein” in maintaining mother-daughter centriole pair cohesion, potentially advancing our comprehension of the pathogenesis of Alström Syndrome.

**Significance statement:** Alström syndrome (AS) is a human ciliopathy that results from mutations in the ALMS1 gene inherited in an autosomal recessive manner. Elucidating the roles of ALMS1 and the underlying molecular mechanisms of AS is of paramount importance. In this study, using *Drosophila* model, we discovered that ALMS1 is localized at the proximal centrioles, and is crucial for the proper centriole engagement, spindle polarity, cell division in spermatocytes. Our findings reveal the new role of ALSM1 in maintaining centriole engagement, and suggest that non-ciliary function of ALMS1 may contribute to the pathogenesis of Alström Syndrome, warranting further investigation.

## Introduction

The centrosome is the major microtubule-organizing center (MTOC) of animal cells, and it typically contains a tightly connected (engaged) centriole pair (consisting of one old mother centriole and one young daughter centriole) arranged in an orthogonal configuration and surrounded by pericentriolar materials (PCM). Through its MTOC activity, the centrosome plays a crucial role in the process of cell division by organizing spindles. Each cell cycle begins with a single centrosome that contains two centrioles. During the S phase, these two centrioles duplicate to form two centrosomes, which are connected by a flexible intercentrosomal linker and act as a single MTOC until late G2 phase. During the G2/M transition, the intercentrosomal linker is broken, and the two centrosomes separate and gradually migrate to opposite poles of the cell to organize a bipolar spindle. At the end of mitosis, the mother-daughter centriole pair is disengaged to allow centrioles to replicate and form two pairs in the next S phase (1, 2). It is believed that daughter centriole remains engaged to its mother centriole until late mitosis to preventing centriole over duplication during the cell cycle, and that disengagement of centrioles at the end of mitosis/meiosis licenses centrioles for duplication in the next cell cycle (3, 4). The timing of centriole duplication, centrosome separation and centriole disengagement are strictly controlled within a cell cycle to ensure proper centriole number, spindle polarity, genome stability, and cytokinesis(5, 6).

The intercentrosomal linker that emanates from the proximal end of the parent centrioles, is composed of proteins associated with proximal centrioles, such as C-Nap1/Cep250 (7), filamentous protein Rootletin (8), and filament modulator Cep68 (9, 10). Centlein (11) and LRRC45 (12) at the proximal ends of centrioles have also been implicated in linker formation. However, it is still unclear how the mother-daughter centriole pair is tightly engaged. One plausible model is that centrioles are held together by “glue” proteins (13), but the identity of these proteins remains elusive. While the cohesion complex has been suggested as a potential candidate to hold mother-daughter centrioles together in mammalian cells (14, 15), its role in maintaining centriole engagement may not be conserved in *Drosophila* (16). Therefore, further investigation is necessary to identify the unknown “glue” proteins responsible for maintaining centriole engagement.

The PCM surrounding centrioles has also been shown to contribute to centriole engagement and centrosome cohesion in both mammalian cells and *Drosophila*. Studies have demonstrated that the removal of major PCM components such as Cnn/Cep215/Cdk5Rap2 (9, 13, 17) or PCNT/Plp (13, 18) resulted in premature disengaged/unpaired centrioles. Inhibition of PCNT cleavage strongly prevents both centrosome separation and centriole disengagement (13). Furthermore, Plk1 has been found to play a critical role in centriole disengagement or centrosome cohesion (19–21). Phosphorylation mediated by Plk1 activates the degradation of PCM or linker proteins via proteasomes or Separase (13, 21–23).

Centrioles not only forms the core of centrosomes, but also serve as templates for cilia formation. Mutations in many centriole proteins lead to ciliopathies (5, 6). Alström syndrome (AS) is a rare autosomal recessive human ciliopathy, resulted from mutations in human ALMS1, a protein localized at the proximal ends of centrioles (24–27). Although ALMS1 is known to be required for ciliogenesis in mammals (28–30), its specific molecular functions in centrosome and cilia remain to be elucidated. In this study, we reported that *Drosophila* Alms1 is specifically required for giant centriole engagement in spermatocytes. Furthermore, we revealed that Alms1 acts downstream of Sas-6 to maintain centriole engagement, and PCM and centriole size have antagonistic roles in maintaining centriole engagement in *alms1* mutant flies. We propose that Alms1 is a glue protein that localizes to the proximal centriole to maintain mother-daughter centriole engagement.

## Results

### *Drosophila* ALMS1 is required for maintaining meiotic centriole engagement in spermatocytes

As previously reported (24), *Drosophila* has two homologs of *Alms1* gene, CG12179 (*Alms1a*) and CG12184 (*Alms1b*), which are juxtaposed to each other on the X chromosome (Fig. 1A). While Alms1a shares a high degree of sequence similarity with Alms1b, it contains a 300-amino acid insertion at its N-terminal region (Supplementary Fig. 1A). Since Alms1a and Alms1b are both localizes to the basal body of spermatocyte cilia (24) and sensory cilia (Supplementary Fig. 1B), it is possible that they have redundant roles in *Drosophila* ciliogenesis and spermatogenesis. To explore the function of ALMS1 using *Drosophila* model, we generated a null allele of *alms1* mutant, *alms1^2^*, with simultaneously large fragment deletion in both *Ams1a* and *Alms1b* genes by using two guide RNAs (gRNAs) targeting *Alms1a* and *Alms1b*, respectively (Fig. 1A, Supplementary Fig. 2).

**Figure 1.**
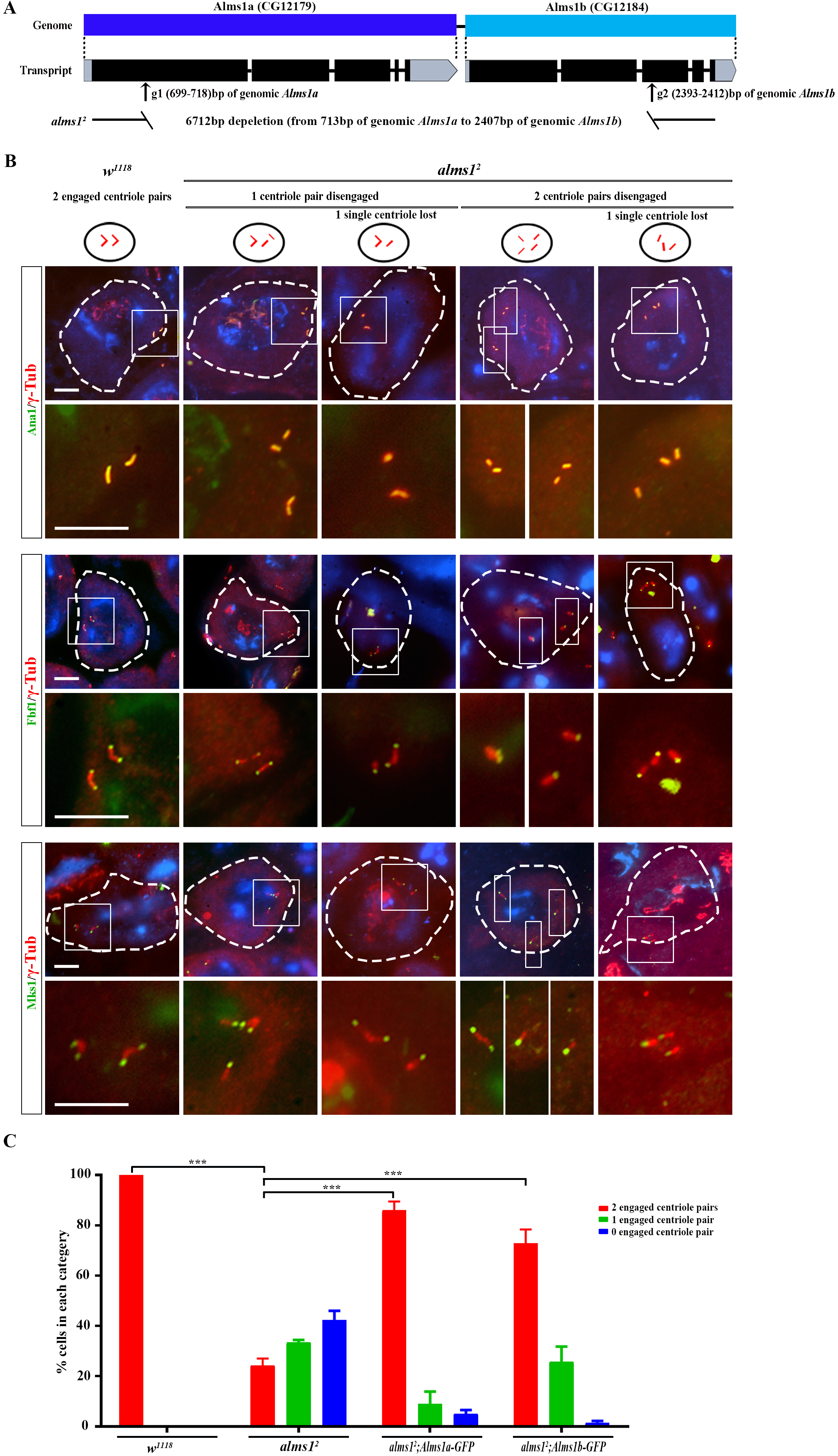
Alms1 is required for maintaining meiotic centriole engagement in *Drosophila* spermatocytes. A): Construction of *alms1* mutants. Graphic illustration of two *Drosophila Alms1* gene- *Alms1a* (CG12179) and *Alms1b* (CG12184). gRNA2 (699-718bp) in *Alms1a* and gRNA1 (2,393-2,412bp) in *Alms1b* were used for generating *alms1* mutant fly. The *alms1^2^* mutant with a large fragment deletion of 6,712 bp was obtained. F-primer (located at 398-422bp of genomic *Alms1a*) and R-primer (located at 2,734-2,758bp of genomic *Alms1b*) were designed for PCR genotyping of *alms1^2^* mutant. B): *alms1^2^*mutant flies showed severe centriole engagement defects in meiotic spermatocytes, displaying modified centriole conformation of 2 engaged centriole pairs, 1 engaged centriole pair (1 engaged centriole pair + 2 singlets; 1 engaged centriole pair + 1 singlet), or 0 engaged centriole pair (4 singlets; 3 singlets). Dotted white circles outline spermatocyte cells. γ-Tub: centrosome marker; Ana1: centriolar protein, Fbf1: transition fiber protein; Mks1: transition zone protein. DAPI labels nuclei. Bars: 10 μm. C): Over-expression of Alms1a-GFP or Alms1b-GFP can well rescue the premature centriole disengagement in *alms1^2^* spermatocytes. Statistical significance was determined using a two-tailed Student’s t-test. ns, P > 0.05; *, P ≤ 0.05; **, P ≤ 0.01; ***, P ≤ 0.001. Error bars represent SEM.

Flies with ciliary defects are often uncoordinated in walking and flying, and exhibit defects in sensory behaviors (31–33). However, our *alms1^2^* mutant flies are viable and exhibit normal walking and flying abilities. Additionally, they do not display any obvious defects in cilia-related sensory behaviors, such as hearing, touch response and olfactory (Supplementary Fig. 3A-D), suggesting that Alms1 is dispensable for sensory cilia related neuronal sensation in *Drosophila*. Previous studies using RNA interference (RNAi) approaches have suggested that Alms1a is involved in centriole duplication in male germline stem cells in *Drosophila* (24). However, in our *alms1^2^* mutants, we didn’t observe any significant loss of centrosome in spermatogonia (Supplementary Fig. 3E).

Interestingly, we observed an intriguing novel phenomenon in *alms1^2^*KO mutants that the mother-daughter centriole pairs in meiotic prophase I spermatocytes were frequently disengaged (Fig. 1B, C). In WT meiotic I spermatocytes, there are always two giant “V-shaped” engaged centriole pairs. However, 1 centriole pair or both 2 centriole pairs were frequently disengaged in meiotic I spermatocytes of *alms1^2^* mutants (Fig. 1B, C). It should be noted that these prematurely disengaged singlet in *alms1^2^* spermatocytes was subject to loss, possibly due to the centriole instability induced by this premature centriole disengagement. Remarkably, either *Alms1a* or *Alms1b* gene expression is able to rescue such centriole disengagement defect in *alms1^2^*mutants (Fig. 1C), supporting that *Drosophila* Alms1 is responsible for centriole engagement during meiosis in *Drosophila* spermatocytes.

### Abnormal male meiotic division in *alms1* mutants

Maintaining the engagement of mother-daughter centriole pairs before telophase is critical for controlling centrosome number and ensuring proper cell division. Premature disengagement of centriole pairs could lead to abnormal spindle organization and incorrect cell division (15, 17, 34–39). In fact, multipolar spindles and chromosome mis-segregation were observed during meiosis in spermatocytes of our *alms1^2^* KO mutants (Fig. 2A).

**Figure 2.**
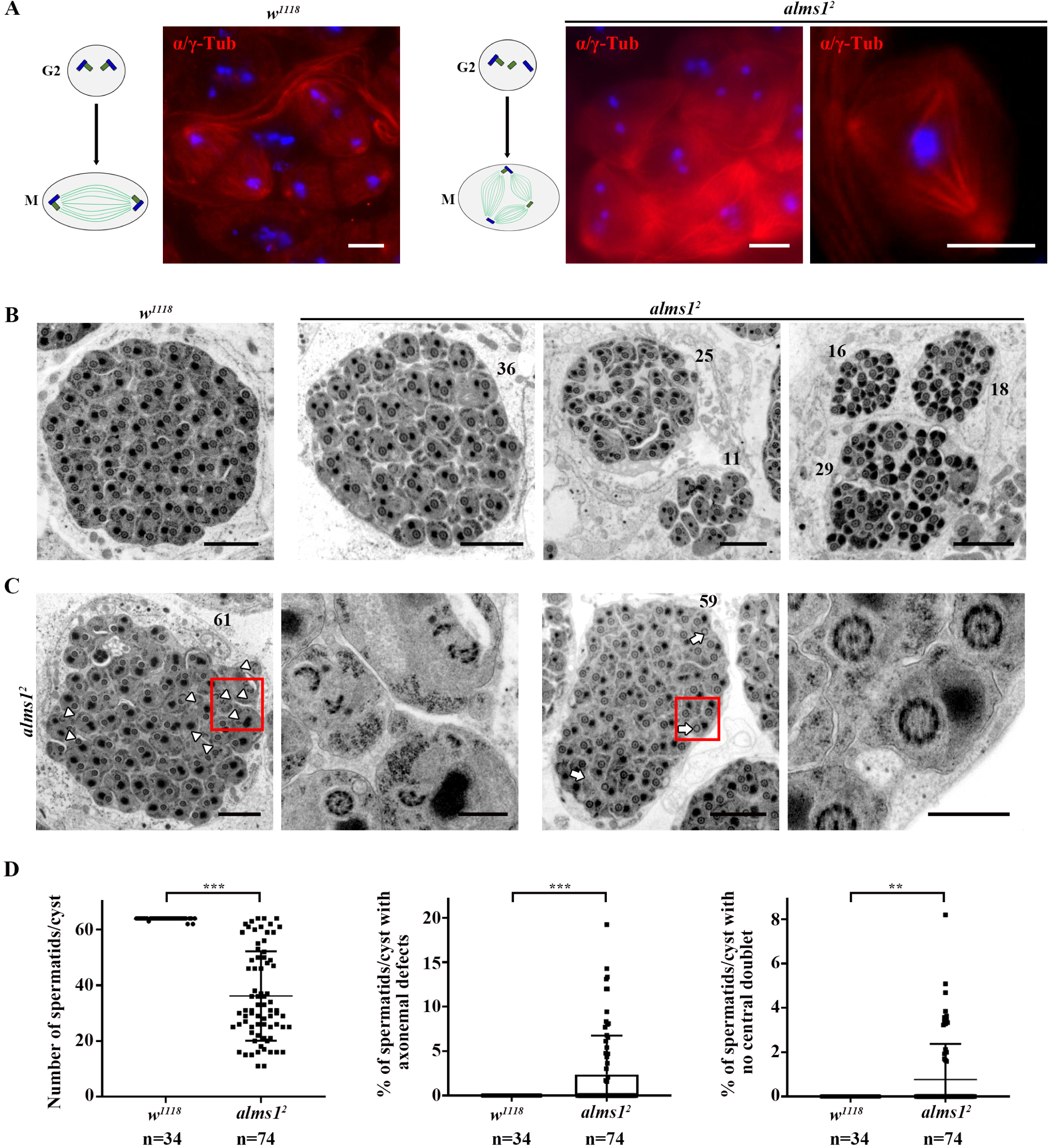
Abnormal spermatogenesis in *alms1* mutant flies. A): In the metaphase of meiosis, bipolar spindles were observed in control spermatocytes (left), whereas, multi-polar spindles were observed in *alms1^2^*spermatocytes (right). Spindles are labeled by α-Tub/γ-Tub; DAPI labels nuclei. B): Alms1 is required for *Drosophila* spermatogenesis. Compared with wild-type spermatid cyst which contains 64 spermatids, the number of spermatids were significantly reduced *in alms1^2^* cysts. C): The flagella in *alms1^2^* spermatids exhibit broken axonemes (white arrow heads) and a lack of central pair microtubules (white arrows). Quantification data was shown in D. Bars: 10 μm (A); 2 μm (B, the 1^st^ row and 3^rd^ row of C); 0.5 μm (the 2^nd^ row and 4^th^ row of C). Statistical significance was determined using a two-tailed Student’s t test. Error bars represent SD. ns, P > 0.05; *, P ≤ 0.05; **, P ≤ 0.01; ***, P ≤ 0.001.

To further assess the defects in *Drosophila* spermatogenesis caused by *Alms1* deletion, we performed transmission electron microscopy (TEM) analysis on spermatid cysts. Our results revealed a significantly reduced number of spermatids per cyst in almost all cysts from our *alms1^2^* mutants (Fig. 2B-D), compared to control *w^1118^* testes where all cysts contain 64 spermatids. This reduction may have resulted from abnormalities in cell division, leading to cell death. We also observed that a fraction of flagella in *alms1^2^* mutants had broken outer-doublet microtubules (Fig. 2C and D), or lacked a central pair of two singlet microtubules (Fig. 2C and D), indicating an important role of Alms1 in *Drosophila* sperm flagella formation. Consistent with this, male fertility is significantly impaired in our *alms1^2^* mutants (Supplementary Fig. 3F). Collectively, our findings highlight the involvement of *Drosophila* Alms1 in spermatogenesis, sperm flagella formation and male fertility, providing valuable insights for further investigation into the mechanisms underlying reduced fertility in male AS patients (40).

### Alms1 functions downstream of Sas-6 to engage centriole pairs

Consistent with previous reports (24), we confirmed that Alms1a localizes to the proximal part of basal bodies, whereas Alms1b is restricted to the proximal end in *Drosophila* spermatocytes (Fig. 3A). Disengaged centriole pair in *Drosophila* spermatocytes was reported previously in *sas-6* mutants (37, 41). Sas-6 is a cartwheel protein that localizes to the proximal end of centriole and is essential for the establishment of centriole’s nine-fold symmetry (37, 42). In addition to premature centriole disengagement, *sas-6* mutants also have flagella that lack central microtubules (37). Considering their comparable subcellular localization and mutant phenotypes, we wonder if there might be a potential relation between Sas-6 and Alms1 in centriole engagement. To answer this question, we first examined their mutual localization dependencies. We observed that either Alms1a or Alms1b is frequently missing from the proximal centrioles of *sas-6* mutant spermatocytes, but the localization of Sas-6 appears to be unaffected in *alms1^2^* mutant spermatocytes (Fig. 3B and C), suggesting that Sas-6 is required for recruiting Alms1 to the proximal centrioles. Next, we investigated their genetic relationships by analyzing the centriole disengagement defects in *alms1*; *sas-6* double mutants, and found that the premature centriole engagement defects in *Drosophila* spermatocytes were not exacerbated by the double mutation. In fact, *alms1*; *sas-6* double mutants showed a similar defect to *sas-6* single mutant, with more severe centriole disengagement defects than those seen in *alms1* single mutants (Fig. 3D). Notably, we observed that overexpression of Sas-6 can partially rescue the premature centriole disengagement defect in *alms1^2^*mutant spermatocytes (Fig. 3E), but conversely, introduction of Alms1a or Alms1b had no impact on the centriole engagement defects in *sas-6* mutants (Fig. 3F). All these results suggest that Sas-6 and Alms1 act in a same pathway, with Sas-6 function upstream of Alms1 to control the centriole engagement of *Drosophila* spermatocytes.

**Figure 3.**
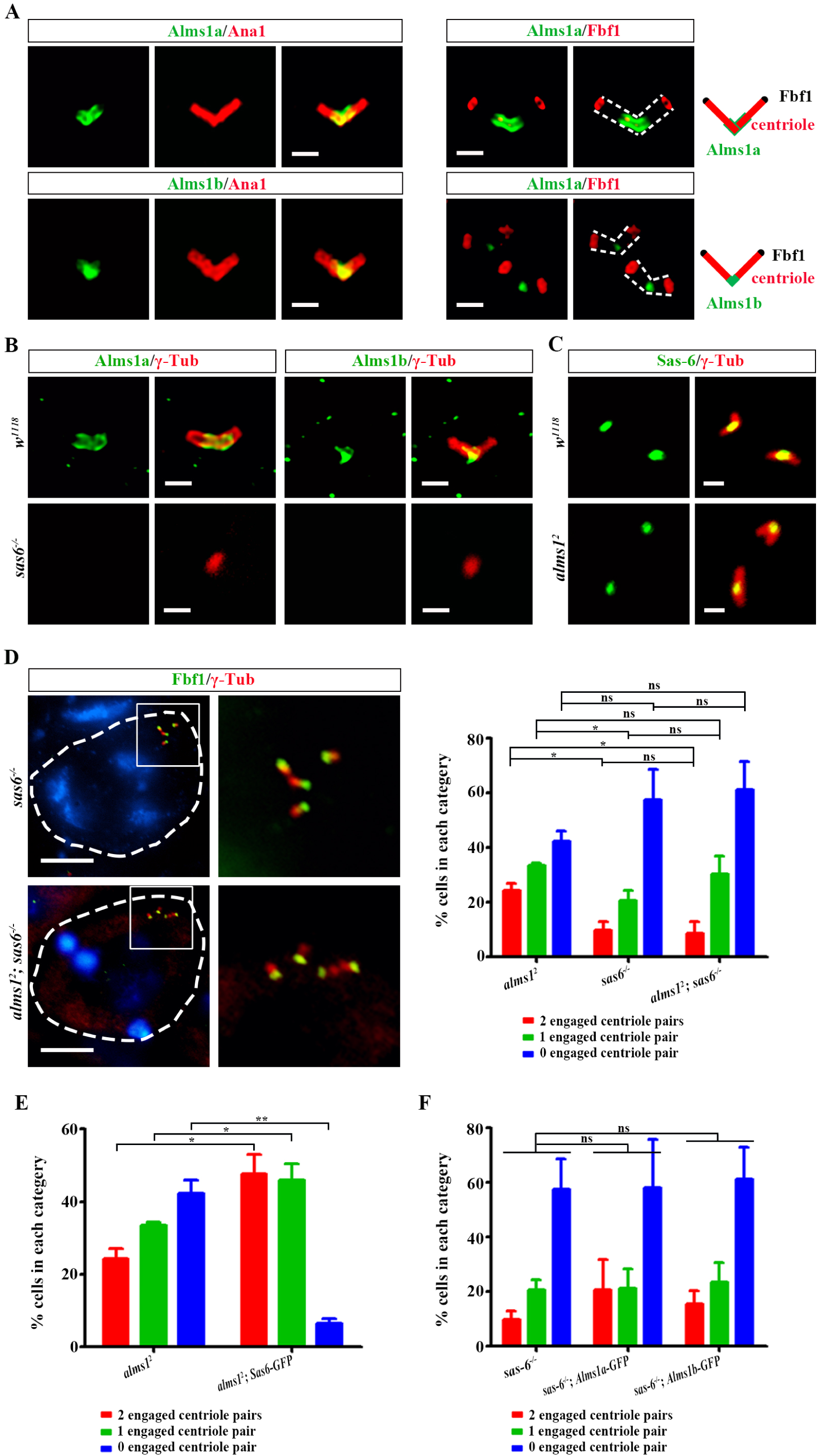
Alms1 functions downstream of Sas-6 to engage centriole pairs. A): *Drosophila* Alms1a and Alms1b were localized to the proximal region of centrioles during spermatogenesis. Green: Alms1a, Alms1b; γ-Tub: centrosome marker; Ana1: centriolar protein; Fbf1: transition fiber protein. Bars: 2 μm. The centrosome is marked with dotted lines. B): Both Alms1a and Alms1b were lost from the proximal of spermatocyte centrioles of *sas-6* mutant (green: Alms1; red: γ-Tub). C): Alms1 is dispensable for Sas-6 localization. D): *alms1^2^*; *sas-6* double mutants showed similar premature centriole disengagement defect as *sas-6* single mutants, but more severe than *alms1^2^* single mutants. Bars: 10 μm. E): Overexpressed Sas-6 partially rescued premature centriole disengagement in *alms1^2^* spermatocytes of *Drosophila*. F): Introduction of *Drosophila* Alms1a-GFP or Alms1b-GFP could not prevent early centriole disengagement defects in *sas-6* spermatocytes. Statistical significance was determined using a two-tailed Student’s t test. ns, P > 0.05; *, P ≤ 0.05; **, P ≤ 0.01; ***, P ≤ 0.001. Error bars represent SEM.

### The conserved ALMS motif is required, but not essential for centriole engagement

Similar to mammal Alms1, there is an evolutionarily conserved ALMS motif at the C terminus of both Alms1a and Alms1b (Fig. 4A, Supplementary Fig. 1A). In mammals, ALMS motif is required, but not essential, for centrosome targeting of Alms1 (26). To determine the role of ALMS motif in basal body targeting of *Drosophila* Alms1, we deleted the ALMS motif in Alms1a and Alms1b, and generated Alms1a^ΔCT^-GFP and Alms1b^ΔCT^-GFP transgenes to check their subcellular localization. Alms1a-GFP localizes to the proximal centrioles, however, when examining Alms1a^ΔCT^-GFP, we found that although it can still target to the basal bodies, the intensity of signal is notably reduced (Fig. 4B). On the other hand, Alms1b-GFP is mainly localized to the proximal end of centrioles, whereas nearly 50% of Alms1b^ΔCT^-GFP displays abnormal localization pattern. These abnormalities include irregular distribution along the centriole, weak signals, or even a complete loss of the centriole localization signal (Fig. 4C).

**Figure 4.**
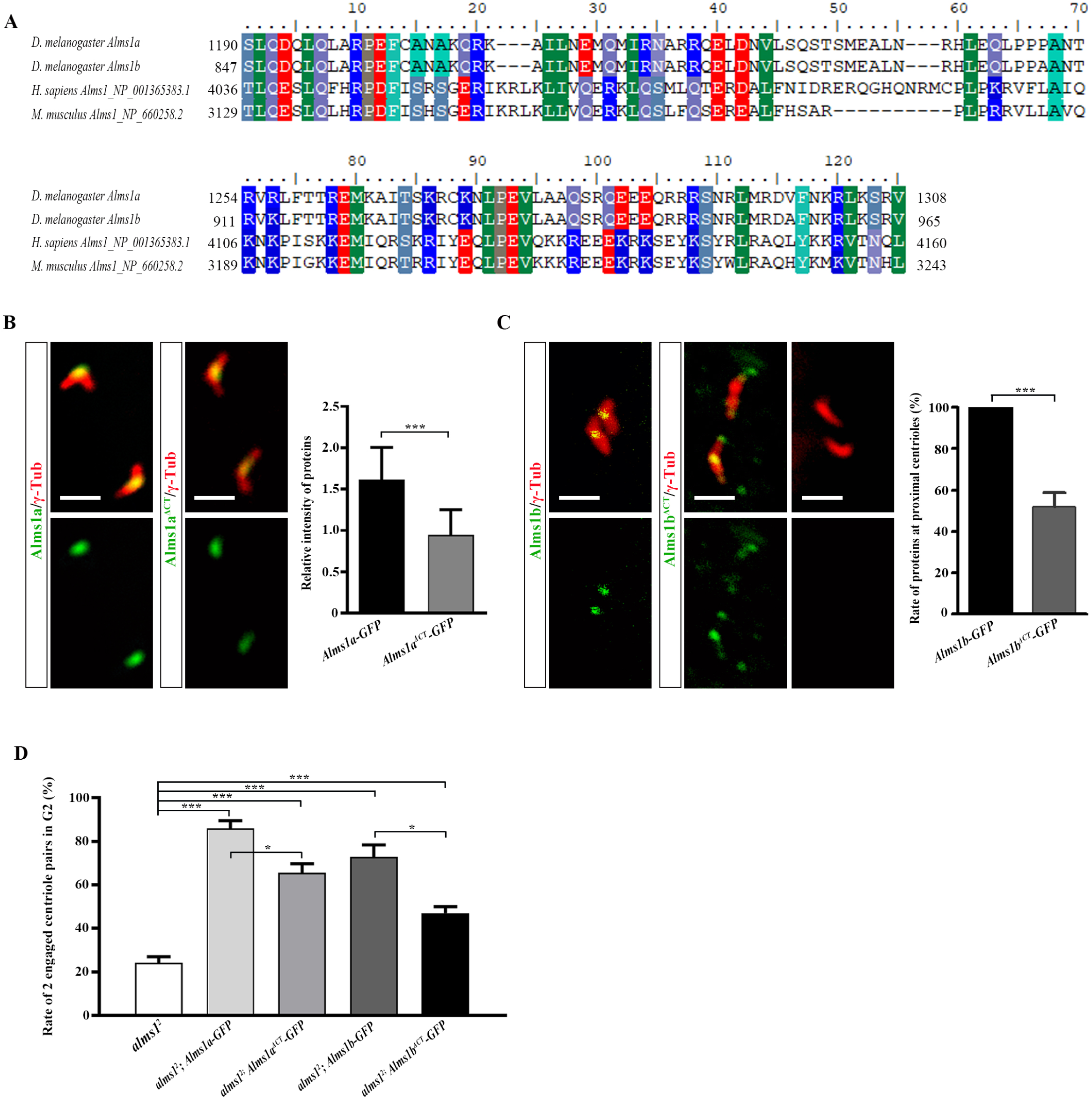
ALMS motif is required but not essential for its centrosomal localization and centriole engagement. A): Alignment of ALMS motif (125aa) of *Drosophila* Alms1, *H. sapiens* Alms1 (NP_001365383.1) and *M. musculus* Alms1 (NP_660258.2). B and C): ALMS motif in *Drosophila* Alms1a and Alms1b is required for its localization at proximal centrioles. Compared to Alms1a-GFP, Alms1a^ΔCT^-GFP was still able to target to the basal bodies, but the intensity of signal is notably reduced (B); nearly 50% of Alms1b^ΔCT^-GFP displays abnormal localization pattern, including irregular distribution along the centriole, weak signals, or even a complete loss of the centriole localization signal (C). D): Alms1a^ΔCT^-GFP or Alms1b^ΔCT^-GFP were able to rescue the centriole disengagement defect in *alms1^2^* mutants, but their efficiency was significantly lower when compared to the intact Alms1 protein. Statistical significance was determined using a two-tailed Student’s t test. ns, P > 0.05; *, P ≤ 0.05; **, P ≤ 0.01; ***, P ≤ 0.001. Error bars represent SEM.

Next, we evaluated the function of ALMS motif by assessing the abilities of Alms1a^ΔCT^ or Alms1b^ΔCT^ to rescue the centriole disengagement defect in *alms1^2^*mutant spermatocytes. We found that both Alms1a^ΔCT^ and Alms1b^ΔCT^ were capable of rescuing the centriole disengagement defect in *alms1^2^* mutants, although they were significantly less efficient compared to intact Alms1 proteins (Fig. 4D). Together, our results suggest that while the conserved ALMS motif of *Drosophila* Alms1 is required, it is not essential for the basal body targeting and functions of Alms1.

### Inactivation of Plk1 could not rescue the premature centriole disengagement of *alms1* mutants

Centriole disengagement requires Plk1 activity which promotes degradation of PCM (13, 23, 39, 43, 44). Inactivation of Plk1/Polo blocks centriole disengagement in both mammals and *Drosophila* (19–21). In *Drosophila* testis, when Plk1 activity is inhibited by BI2536, a Plk1/Polo inhibitor, paired centrioles remain engaged after Meiosis II of spermatocytes, which leads to the emergence of two flagella from a round spermatid (Fig. 5A). We wondered whether inhibition of Plk1 could overcome the centriole engagement defect of *alms1^2^* mutants. However, treatment with BI2536 did not improve the premature centriole disengagement defects in *alms1^2^* spermatocytes, and there was no significant increase in the percentage of engaged centriole pairs (Fig. 5B). These results indicate that Alms1 functions in a distinct pathway from PLK1 to maintain centriole engagement, and centriole disengagement caused by Alms1 deletion occurs prior to PLK1-mediated centriole disengagement at the end of meiosis, suggesting that Alms1 may act as a potential “glue protein” to maintain the engagement of mother-daughter centriole pairs throughout the cell cycle in *Drosophila*.

**Figure 5.**
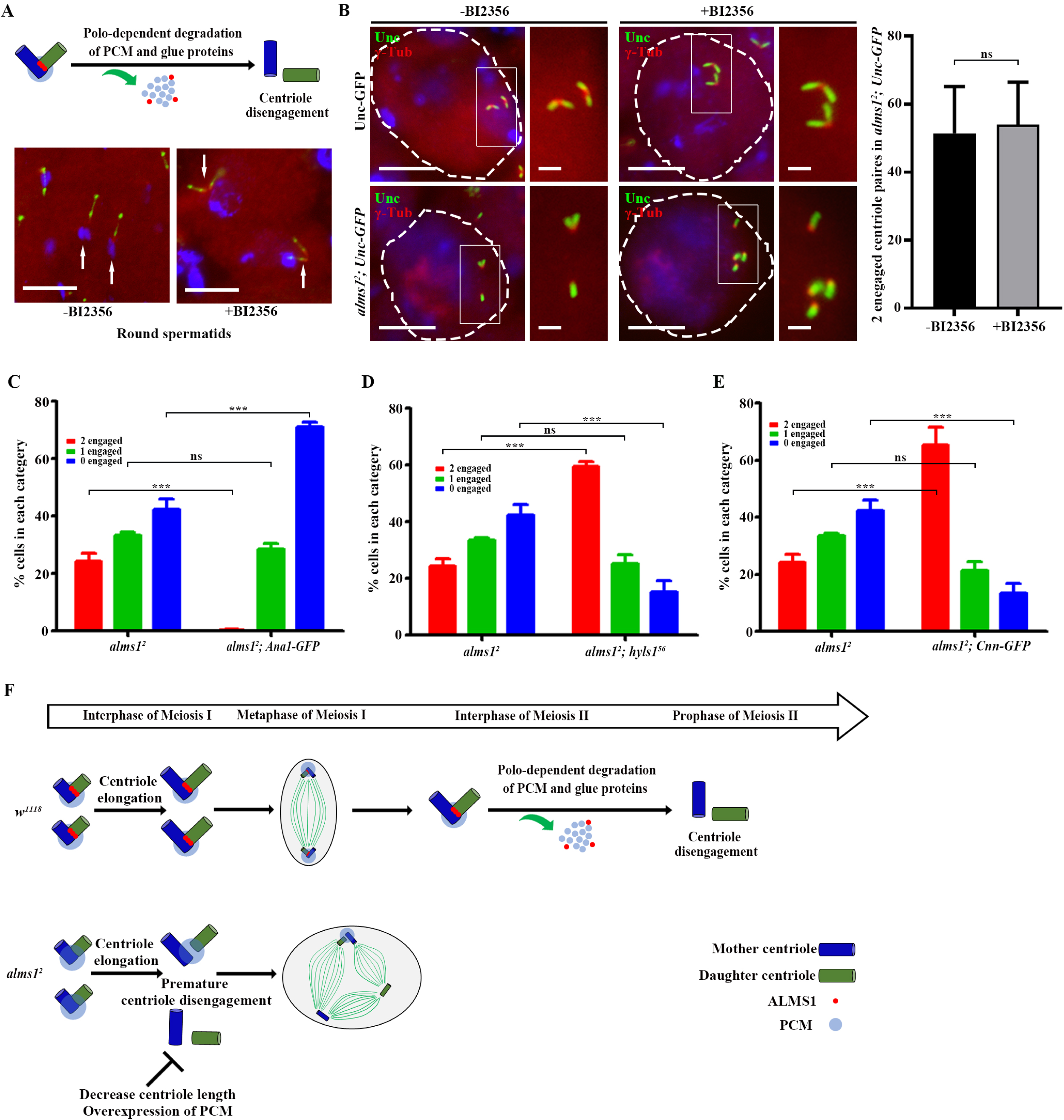
Role of PCM, centriole length and Polo in Alms1-mediated centriole engagement. A): Centriole disengagement in male meiosis is inhibited by Polo inhibitor, BI2536. Engaged centriole pairs still could be observed after meiosis II (white arrows). Unc-GFP labels the flagella; γ-Tub (red) indicates the basal body. B): Inactivation Polo by BI2536 could not rescue the premature centriole disengagement defect in *alms1^2^* spermatocytes. Bars: 10 μm (A, the 1^st^ and the 3^rd^ row of B); 2 μm (the 2^nd^ and the 4^th^ row of B). C and D): Modifying the length of centriole altered the incidence of centriole disengagement in *alms1^2^* mutants. Longer centrioles by overexpressed Ana1-GFP led to more severe premature centriole disengagement (C); and shorter centrioles by deletion of *Hyls1* rescued premature centriole disengagement defect in *alms1^2^* spermatocytes (D). E): Introduction of PCM protein Cnn-GFP could rescue premature centriole disengagement of *alms1^2^* spermatocytes. Statistical significance was determined using a two-tailed Student’s t test. ns, P > 0.05; *, P ≤ 0.05; **, P ≤ 0.01; ***, P ≤ 0.001. Error bars represent SEM. F): Model for the role of *Drosophila* Alms1 in centriole engagement. Centriole engagement is maintained by the complex display among linkage proteins, PCM and the size of centrioles. In WT, meiotic centriole disengagement occurs in prophase of Meiosis II, where PCM and linkage proteins were removed in a in a Polo-dependent manner. Alms1 deletion results in weaker linkage between large mother and daughter centriole, results in early disengagement in Meiosis I and led to multipolar spermatocyte cells. Increased centriole length worsens the defects, and overexpression of PCM stabilizes the centriole engagement. Blue: mother centriole; green: daughter centriole; red: Alms1, light blue: PCM.

### Antagonistic role of PCM and centriole length in maintaining centriole engagement in *alms1* mutants

The requirement for Alms1 in maintaining the centriole engagement is likely to be specific to spermatocytes, as the cell division appears normal in embryos (Supplementary Fig. 3G). Unlike centrioles in other tissues, centrioles in *Drosophila* spermatocytes are extra elongated, forming a large “V-shaped” centriole pair. We speculate that this tissue-specific difference may be attributed to the size of the centrioles, as larger centriole pairs would require stronger linkage to maintain their engagement. Therefore, we hypothesized that modifying the size of centrioles in *alms1^2^* mutants would affect the incidence of premature centriole disengagement. To test this, we increased the centriole length by overexpressing Ana1-GFP, a positive regulator of centriole assembly (45, 46). As expected, larger centrioles indeed worsened the premature centriole disengagement defect in *alms1^2^* spermatocytes (Fig. 5C). Conversely, we decreased centriole length of *alms1^2^* mutant by deletion of HYLS1, which is known to cause shorter centrioles (47), and found that the premature centriole disengagement defect was significantly restored in *alms1^2^; hyls1^56^* double mutants (Fig. 5D). These results provide strong evidence that the size of centrioles does affect the engagement between mother-daughter centrioles.

It has been reported that the pericentriolar material (PCM), surrounding the mother-daughter centriole pair, helps stabilize/enhance the centriole engagement/cohesion. Previous researches have shown that two critical PCM components, PLP and CNN/Cep215/Cdk5Rap2, are involved in the tight engagement of mother-daughter centriole pairs (9, 13, 17, 23, 39, 43). Building upon these findings, we hypothesized that overexpression of Cnn would be possible to stabilize the centriole engagement in *alms1^2^* mutants. As expected, overexpression of centrosomal Cnn indeed efficiently rescued the premature centriole disengagement defects in *alms1^2^* spermatocytes (Fig. 5E). Therefore, we conclude that the centriole size and PCM integrity antagonistically regulate Alms1-mediated centriole engagement in *Drosophila* (Fig. 5F).

## Discussions

As a disease-causing gene, Alms1 has received great attention in the past few years (Alvarez-Satta et al., 2015; Choudhury et al., 2021; Collin et al., 2012; Hearn, 2019; Maffei et al., 2002; Marshall et al., 2007a; Marshall et al., 2011; Marshall et al., 2015). Despite progress in understanding the biological function of Alms1 (Alvarez-Satta et al., 2015; Hearn, 2019), the molecular function of Alms1 in centriole/cilia is still largely unknown. In this study, we demonstrated that Alms1 is critical for centriole engagement during male meiosis in *Drosophila*. Previously, it has been reported that Alms1 is required for maintaining centrosome cohesion in mammalian cells (Hearn, 2019; Knorz et al., 2010). It is important to note that the structure that engages mother-daughter centriole pair is distinct from the intercentrosomal linker responsible for centrosome cohesion. Nevertheless, it is highly possible that some proteins are shared by these structures at the proximal end of centrioles. As a result, Alms1 could potentially play a role in both centriole engagement and centrosome cohesion. Additionally, proteins in the intercentrosomal linker are closely related to the ciliary rootlet (Bahe et al., 2005; Pagan et al., 2015), a structure implicated in ciliogenesis (Yang et al., 2002). Given this connection, it is possible that Alms1 regulates ciliogenesis by affecting ciliary rootlet. It would be interesting to investigate whether the role of Alms1 in centriole engagement is conserved in mammals, and to explore the role of Alms1 in ciliary rootlet.

Although the maintenance of centriole engagement is critical for cell cycle, the underlying molecular mechanism remains unclear. A plausible model is that glue proteins connect the cartwheel structure at the proximal end of daughter centriole and centriole wall at the proximal of mother centriole. Consistent with this model, studies have revealed the critical role of cartwheel proteins in centriole engagement. In *Drosophila*, deletion of major cartwheel protein Sas-6 or Ana-2 results in premature centriole disengagement (37, 41). Cohesion have been suggested as a glue protein in mammals, but its function in *Drosophila* is not conserved (16). Here, we discovered that the centriole proximal protein Alms1 plays a role in centriole engagement during male meiosis in *Drosophila*. Our research demonstrated that Alms1 functions in a same pathway as Sas-6, but downstream, to regulate centriole engagement. It has been reported that Alms1 directly interacts with Plk4, therefore, Alms1 may interact with the cartwheel via Plk4 (24). To fully understand the molecular basis of Alms1’s role in centriole engagement, a detailed investigation of its molecular interactions with the mother centriole wall is necessary.

We have shown that modifying the size of the centriole and the pericentriolar material (PCM) can affect the incidence of centriole engagement in the *alms1^2^*mutant, providing compelling evidence that centriole size and PCM play a role in regulating centriole engagement. Previous studies have also highlighted the involvement of PCM in centriole engagement (13, 17, 18, 39, 48). Thus, the maintenance of centriole engagement requires a complex interplay among linkage proteins, PCM, and the size of centrioles.

Several extraciliary functions of Alms1 have been reported. These include cell cycle regulation (49, 50), centrosome duplication (24), apoptosis (51), and endosome recycling (52). Additionally, Alms1 has been suggested to function in transcription (53), DNA damage response (54, 55), and other process. Given that defects caused by abnormal centriole engagement may extend beyond cilia, it would be intriguing to explore the connection between Alms’s extraciliary functions and centriole engagement. Further investigations are needed to determine the potential roles of non-ciliary functions of Alms1 in the pathogenesis of Alström Syndrome.

## Materials and Methods

### Fly stocks

Alms1a-GFP, Alms1b-GFP, Alms1a^ΔCT^-GFP, Alms1b^ΔCT^-GFP and Sas-6-GFP are generated in this study, Ana1-GFP, Unc-GFP, Cnn-GFP, *hyls1^56^* and Mks1-GFP were described previously (47, 56). *w^1118^* were used as wild-type controls. All flies were kept at 25.

For generating transgenes of Alms1a-GFP, Alms1b-GFP, Alms1a^ΔCT^-GFP and Alms1b^ΔCT^-GFP under the control of an ubiquitin promotor, the entire coding region or fragment with no ALMS motif was cloned into the pJFRC14-GFP vector, respectively. To generate an *alms1* mutant via CRISPR/Cas9 system, two guide RNA (gRNAs) targeting to *Alms1a* and *Alms1b* was respectively cloned into a modified Peasy-blunt vector with a pU6.3 promotor, and then together injected into the one-cell stage embryos of Cas9. The *alms1^2^* mutant line was obtained, and the mutation was confirmed through genomic PCR and sequencing. The mutant was backcrossed with *w^1118^* before behaviors analysis.

Primers used:

F primer for Alms1a-GFP / Alms1a^ΔCT^-GFP transgene:
5′- GCAGAATAATCCACCAGATCTATGAGAGCGAAGAGAGGAGC -3′
F primer for Alms1b-GFP / Alms1b^ΔCT^-GFP transgene:
5′- GCAGAATAATCCACCAGATCTATGTTTAAGAAGTTTGGTTCC -3′
R primer for Alms1a-GFP transgene:
5′- TTCTCCTTTACTCATGGATCCTTCGATCTTCTCTAGATGATTAC -3′
R primer for Alms1a^ΔCT^-GFP transgene:
5′ -TTCTCCTTTACTCATGGATCCGCCCGTCGGCGTAAACTGGAT -3′
R primer for Alms1b^ΔCT^-GFP transgene:
5′- TTCTCCTTTACTCATGGATCCCTCACTGCTGCCCGTCGGCGT -3′
gRNA1: 5′- GCCAACACAGCAGCGACCCG -3′
gRNA 2: 5′- ACATAGCTAGCTACTCCCTG -3′
test F primer: 5′- AAGTAACTAAGACGAAACCGAATGA -3′
test R primer: 5′- GCGAATCATCTGCATCTCATTAAGT -3′

### Antibodies

The following primary antibodies were used for immunofluorescence in *Drosophila*: γ-tubulin (1:200, Abcam), α-tubulin (1:200, Sigma-Aldrich), Fbf1 (1:200, generated by our lab), rabbit anti-GFP (1:1000, Abcam), mouse anti-21A6 (1:200, DSHB). The secondary antibodies were Alexa Fluor 488 or 594-conjugated goat anti-mouse IgG or anti-rabbit IgG (1:1,000; Abcam).

### Male fertility assay

*alms1^2^* mutant males and virgin females from *w^1118^* were respectively collected and kept for 3-4 days before crossing. Each *alms1^2^* male was mated to one *w^1118^ virgin* for 4 days, followed by removing the parents. Percentage of fertile *alms1^2^* mutants was recorded. All crosses were performed at 25.

### Negative gravity assay

Adult flies, aged 3-4 days, were transferred into a 20-cm-long glass vial that was divided into four sections for testing. Before each test, flies were gently tapped down to the bottom, and then give 15s to climb/fly, after which they were scored based on the height of their climb: 0-5 cm, 1; 5-10 cm, 2; 10-15 cm, 3; 15-20 cm, 4. After each test, flies were given 1min for recovery. To calculate the climbing index, flies were tested in 10 replicate batches and the average score was recorded. At least 5 groups were tested.

### Testis and antennae staining

Fresh testes of young males, antennae of pupae were dissected in fresh 1 × PBS and immediately transferred to a charged microscope slide. After softly squashed with coverslip, samples were immersed in liquid nitrogen for ∼20s, and then fixed in -20 methanol for 15min and -20 acetone for 15min. After washing 15 min×3 times in 1 × PBS buffer, samples were blocked at room temperature for 1 hour with blocking buffer (1 × PBS + 3% BSA + 0.1% Triton), followed by incubation with proper primary antibodies at 4 overnight. Slides were washed with PBT (1× PBS + 0.1% Triton) and incubated with secondary antibodies at room temperature for 2-3 hours. Nuclei were stained with DAPI (1mg/ml, Invitrogen) for 30 min. Samples were mounted in 50% glycerol for imaging (Nikon, Ti2) or in ProLong Diamond Antifade Mountant with DAPI (Invitrogen) for lightning imaging (Leica, LAS X).

### Embryo whole mount staining

Flies that are 2-3 days old were transferred to a fresh agar plate in egg-collecting chambers. After 8 hours of crossing, adult flies were transferred to a new plate for collection of embryos at nearly the same developmental stage. After 2 hours, adults were removed and embryos were kept at 25 for 2 hours before they were immersed in 5% NaClO for dechorionation. Followed by resuspended in a mixture of heptane and 4% PFA (1:1) for removing vitelline membranes, samples were washed in 1 × PBS 15min× 3 times, and blocked in PBST (PBS + 0.1% BSA + 0.1% Triton) for 1 hour at room temperature before incubation with primary antibodies. Thereafter, embryos were washed 3 times in PBT (PBS + 0.1% Triton), and incubated with secondary antibody for 2-3 hours at room temperature. Nuclei was stained with DAPI (1mg/ml, Invitrogen) for 30 min. After 3×15 min washes in PBS, embryos were mounted and imaged with Nikon Ti-2.

### Transmission electron microscope

Testes were dissected in prechilled 1 × PBS and immediately transferred to 2.5% glutaraldehyde at 4°C for at least 24 hour, then postfixed in 1% OsO4. Samples were washed in 1M PBS three times for 15 min, and dehydrated through alcohol series and propylene oxide. After, testes were embedded in Epon medium. Ultrathin sections were cut with an ultramicrotome (Leica), and examined under a transmission electron microscope (HT-7700, 120KV).

### Polo inhibition assay

As previously described by Riparbelli (20), testes from 5-7 days old pupae were dissected in prechilled PBS and immediately kept in M3 medium containing 100 nm BI2536 for 24h at 24°C. After incubation, the testes were washed in M3 medium for 10 min and then in PBS for 5 min. Following examination was processed as described in “Testis and antennae staining”.

## Acknowledgments

We thank the Core Facility of Drosophila Resource and Technology, CEMCS, CAS for creating mutants and *Drosophila* transgenes. This research was funded by National Natural Science Foundation of China (no. 32070692 and 31871357 to Q. W., and 31802009 to Y. H.) and China Postdoctoral Science Foundation (no. 2020M672884) to Y. H.

## Author contributions

Y. H. and Q. W. designed the study; Y. H. performed experiments; Y. H. and Q. W. wrote the manuscript. Z. W. assisted the experiments. All authors discussed the results and commented on the manuscript.

## Competing Interest Statement

The authors declare no competing or financial interests.

---

**Supplementary Figure 1.**
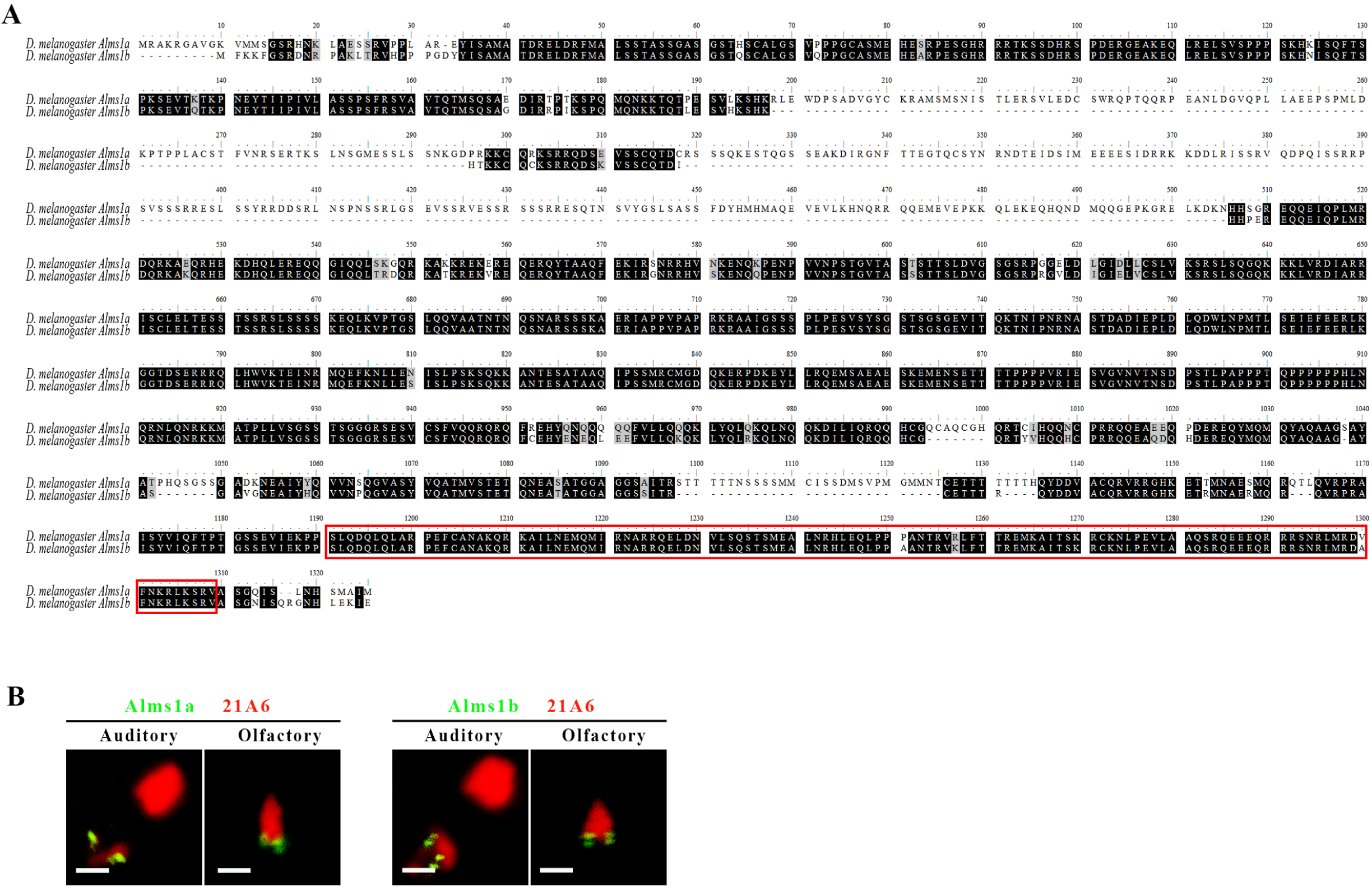
The alignment of *Drosophila* Alms1a and Alms1b, and their localization in sensory cilia. A): The alignment of *Drosophila* Alms1a and Alms1b. Alms1a and Alms1b share high sequence similarity, with the exception of a 300-amino acid insertion in the N-terminal region of Alms1a. The conserved ALMS motif is marked with red rectangle. B): Both Alms1a-GFP and Alms1b-GFP were localized at basal bodies of ChO cilia (Auditory organ at 2^nd^ segment of antennae) and EsO cilia (Olfactory organ at 3^rd^ segment of antennae). 21A6 labels the cilium proximal end of sensory cilia and the extracellular region right below the ciliary dilation of ChO cilia; bars: 5μm.

**Supplementary Figure 2.**
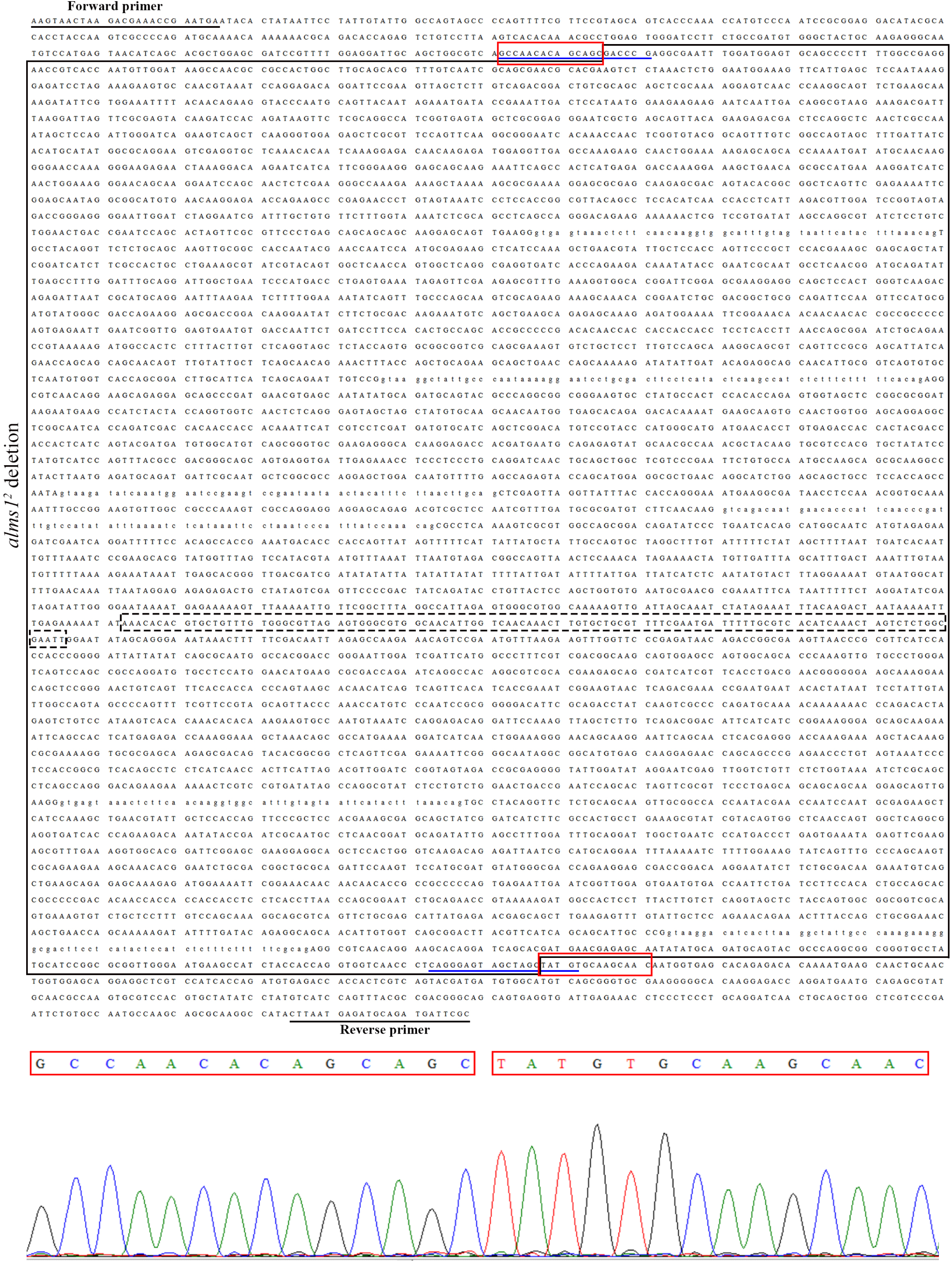
Sequence information on the mutation in *alms1^2^* mutant. A large fragment of 6,712 bp (black frame) was removed in *alms1^2^* mutant. A 113-bp gap between *Alms1a* and *Alms1b* is present and marked with a dotted rectangle. F primer and R primer are marked with black underlines. The locations of two gRNAs used for *alms1* mutant generation are underlined with blue. Red frames label the boundary of *alms1^2^* deletion.

**Supplementary Figure 3.**
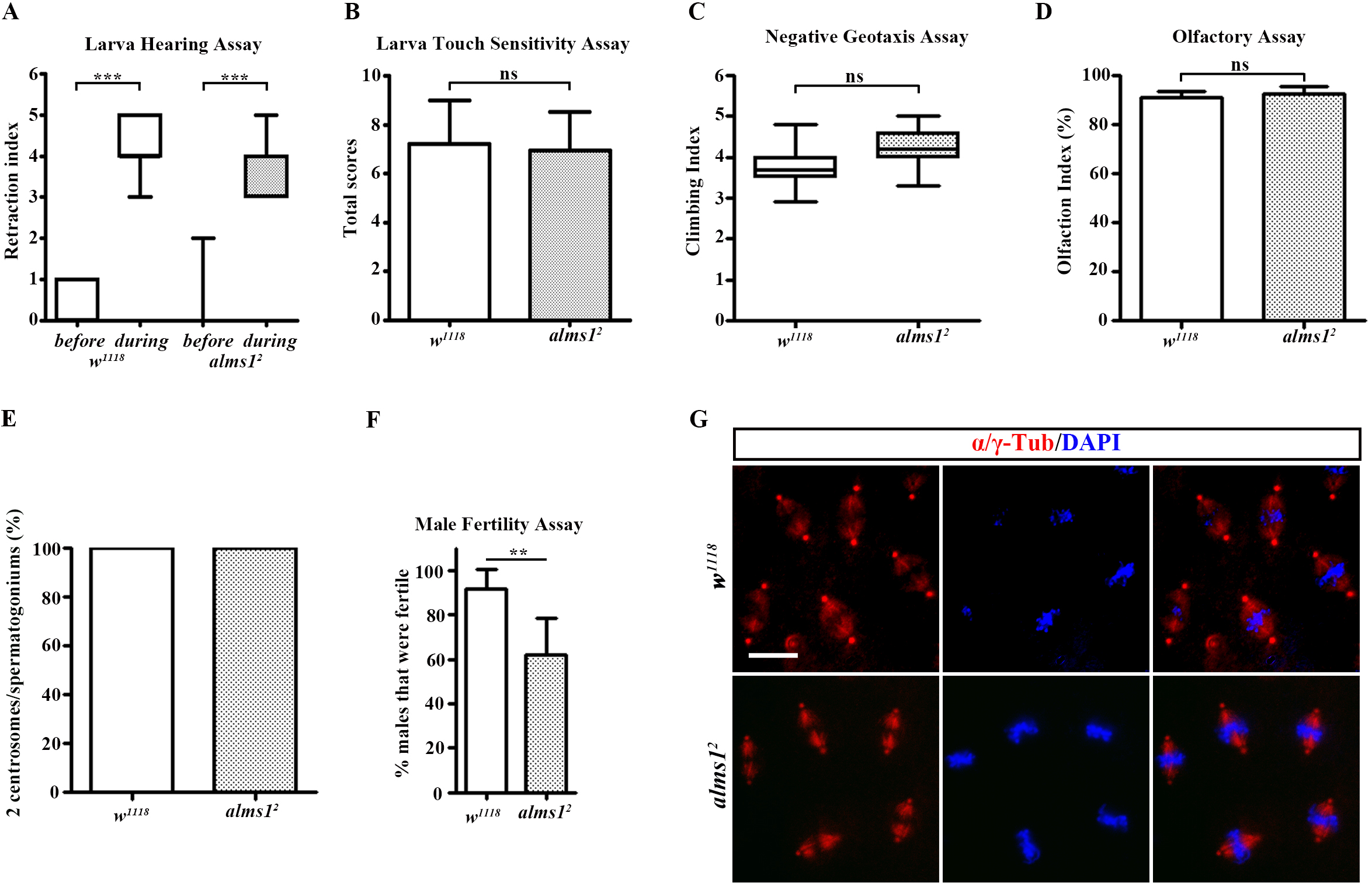
The phenotypes of *Drosophila alms1* mutants. A-D): The sensory behavior assays of *alms1* mutants. *Drosophila* Alms1 was dispensable for hearing capability (A), touch sensitivity (B), negative geotaxis behavior (C) and olfactory responsiveness (D). E): We did not observe any significant loss of centrosome in spermatogonia of *alms1^2^* mutant. F): The *alms1^2^*mutant males exhibited impaired fertility. G): No abnormal spindle organization and cell division were observed in *alms1^2^* mutant embryos. α-Tub: labels spindle; γ-Tub: labels centrosome; DAPI: labels nuclei. Statistical significance was determined using a two-tailed Student’s t test. ns, P > 0.05; *, P ≤ 0.05; **, P ≤ 0.01; ***, P ≤ 0.001. Error bars represent SEM (A-D, F), SD (E). Bar: 10 μm.

## Notes

### Competing Interest Statement

The authors have declared no competing interest.

